# Quantitative measurements of shear stress-dependent adhesion and motion of *Dictyostelium discoideum* cells in a microfluidic device

**DOI:** 10.1101/2024.10.12.618005

**Authors:** Sepideh Fakhari, Clémence Belleannée, Steve J. Charrette, Jesse Greener

## Abstract

Shear stress plays a crucial role in modulating cell adhesion and signaling. We present a microfluidic shear stress generator used to investigate the adhesion dynamics of *Dictyostelium discoideum*, an amoeba cell model organism with well-characterized adhesion properties. We applied shear stress and tracked cell adhesion, motility, and detachment using time-lapse videomicroscopy. In the precise shear conditions generated on-chip, our results show cell migration patterns influenced by shear stress, with cells displaying an adaptive response to shear forces as they alter their adhesion and motility behavior in reaction to shear stress. Additionally, we observed that DH1-10 wild-type *D. discoideum* cells exhibit stronger adhesion and resistance to shear-induced detachment compared to *phg2* adhesion-defective mutant cells and also highlighted the influence of initial cell density on detachment behavior.

## Introduction

Shear stress is an external mechanical force that can influence cell adhesion. This fundamental biological process involves the interaction with and attachment of cells to various surfaces, processes that are usually mediated through specialized protein complexes.^1^ The mechanical aspects of cell adhesion are under intensive ongoing theoretical and experimental investigation^2,3^ due to their crucial role in the behavior of many microorganisms.

Dictyostelium discoideum, a social amoeba that naturally inhabits soil, has been extensively studied for cell adhesion, motility, and development, and other biological processes.^4^ *D. discoideum* undergoes two primary life stages: the first is a unicellular vegetative stage, and the second involves multicellular development. In the first stage, the cells employ phagocytosis for nutrition. This process involves extending pseudopods—temporary cytoplasmic projections—used to engulf and internalize prey.^5^ Successful phagocytosis depends on the amoeba’s ability to adhere to surfaces and move effectively, even under shear stress conditions. When the food source is exhausted, *D. discoideum* cells switch to the second life stage, which starts with the accumulation of hundreds of thousands of cells into dense colonies (pre-aggregation phase) before aggregating into three-dimensional structures and undergo significant changes via gene expression leading to the formation of mature fruiting bodies bearing spores.^6^ During the pre-aggregation phase, the cell density increases beyond 100 cells mm^−2^, reaching at least 1000 cells mm^−2^.^7^

Previous research on *D. discoideum* cell adhesion under shear stress has elucidated several critical mechanisms that govern cellular adhesion and detachment. For example, the number of adhered cells exhibit a sigmoidal decrease with increasing shear stress, suggesting normally distributed adhesion across the cell population.^8^ The initial reversible adhesion is typically followed by a stabilization phase, which highlights the importance of temporal factors in adhesion specificity.^9^ Adhesion force measurements have revealed the role of surface hydrophobicity,^10^ and the presence of biomolecules at the attachment surface.^11^ Other studies have focused on the role of the hydrodynamic environment in cell motility and directionality and determined that shear flow acts as a mechanical cue that influences cellular behavior.^12^

Most studies of cell shear stress adhesion, including those on *D. discoideum*, are conducted using macroscale setups, which involve application of external forces originating from spinning disks,^8^ centrifugation, ^9,13^ or flow chambers (radial^14^ or rectangular^11,15^). However, these conventional methods have limitations, including low throughput, complex assembly, restricted range and limited precision in application of detachment forces, often due in part to non-uniform velocity distributions and turbulence.

Microfluidic devices have become accepted tools for precise measurements pertaining to microorganisms and their multicellular constructs.^16–18^ The low material consumption and high surface-area-to-volume ratios in microchannels can significantly decrease experimental costs and increase cell-surface interactions. Most relevant for the present work, the specific ability to accurately manipulate and control fluids at the micro-/nanoscale offers the potential for detailed studies into cell adhesion and motility due to the combination of highly controllable shear forces and the compatibility of transparent planar microchannels with high-quality microscopy.^19^ Various microfluidic setups have been used to study the effect of shear forces applied to different cell types, including, but not limited to, neutrophils,^20–23^ endothelial cells,^19,24–26^ bacterial cells,^27–31^ and, as we have recently shown, their biofilms.^32–36^

Microfluidic studies of *D. discoideum* have included single-cell studies^37,38^ as well as chemotaxis studies.^39–42^ Other studies used high-resolution time-lapse microscopy to monitor the real-time interactions between *D. discoideum* and bacterial pathogens.^38^ Though less frequent, microfluidic adhesion studies on *D. discoideum* have also been demonstrated. For example, tapered channels in which shear stress was correlated to downstream position (based on the corresponding local channel width) were used to measure detachment kinetics and shear stress-dependent motion under different applied shear stresses.^43^ The results noted a lognormal distribution of the threshold stresses for detachments and first-order kinetics. However, while innovative, the tapered channel design also could also result in artifacts from wall effects due to the coupling of applied shear forces with changes to position in cell motility studies. Tarantola et al. employed a microfluidic device to study surface adhesion of vegetative *D. discoideum*, in which they reported a ten-fold decrease in substratum adhesion.^44^ The highly engineered devices included several channel branch points which admitted liquid to various attachment chambers. Based on the complex interconnected flow paths and chamber sizes, different shear stresses were developed in chambers.^45^ Drawbacks included non-obvious flow properties due to the multitude of branchpoints and complex channel structures. Advantages included large rectangular cell-adhesion chambers in which constant sheer stresses could be applied while minimizing the confounding influence of cell-wall interactions. Although it was not exploited in the study by Tarantola et al., the large cell-adhesion chambers have the potential for motility studies in parallel with adhesion studies.

In this study, we present a microfluidic shear stress generator coupled with time-lapse video microscopy. This device was used to study both the motility and adhesion of native cells. We designed the system to prioritize precision and reliability, with a focus on controlled single-condition experiments rather than automated or high-throughput methods. By using the same chamber and sequentially applying different shear stresses via flow rate adjustments, this approach allowed for precise real-time observation of cell detachment. Our simplified yet effective system enabled accurate quantification of shear stress impacts and initial cell density measurements. Using this platform, we analyzed the detachment and movement of the wild-type *D. discoideum* strain (DH1-10) and a mutant strain (*phg2*). By comparing the behavior of these cells, we validated the ability of our microfluidic shear stress generator to quantitatively assess cell adhesion and investigate the role of cell type and density.

## Materials and Methods

### Cell culture

*D. discoideum* DH1-10^46^ and *phg2* cells^47^ were used in this study. The *phg2* gene encodes the Phg2 protein, a putative serine/threonine kinase essential for phagocytosis, and cell adhesion and mutations in the *phg2* gene impair cell adhesion,^47–49^ which also emphasizes the significance of *phg2* in coordinating adhesion-related signaling pathways and actin cytoskeleton reorganization.^50^ As previously described,^46^ these cells were cultured at 21°C in an HL5 medium supplemented with 15 μg mL^−1^ tetracycline, following the method outlined by Mercanti et al.^51^ The cells were subcultured twice per week in a fresh medium to prevent them from reaching confluence.

### Device fabrication

A microfluidic device with a total volume of approximately 2 µL was designed using standard photolithography techniques,^52^ including CAD software, and a photomask was fabricated by photoplotter (FPS25000, Fortex Engineering Ltd., UK) for use in photolithography. A mold for casting the device was created from a single dry photoresist film (SY300 film, Fortex Engineering Ltd., UK), which was then laminated onto a glass slide (12–550 C, Fisher Scientific, Canada) using a lamination system (FL-0304-01, Fortex Engineering Ltd., UK). The height of the laminated photoresist film determined the final channel height (50 μm). The shadow mask and the photoresist-coated glass slide were exposed to UV light in a UV exposure system (AY-315, Fortex Engineering Ltd., UK) to selectively crosslink portions of the photoresist. Subsequent immersion in developer and rinse baths (SY300 Developer/Rinse, Fortex Engineering Ltd., UK) removed the uncrosslinked portions of the photoresist, resulting in formation of the final mold. A polydimethylsiloxane (PDMS) and crosslinker solution were mixed at a 10:1 ratio, cast on top of the mold, and cured at 70°C overnight. Two inlet holes and one outlet hole were punched into the PDMS at each channel termination point. The two inlets were included in the design for the administration of liquid without the need to disconnect the system during multiple switches between culture media flow and cell flow, thereby preventing the introduction of air bubbles. Air plasma activation (PCD001, Harrick Plasma, USA) was used to bond and seal the PDMS device onto a glass slide. The bonded device underwent brief annealing at 70°C to enhance the bond strength. The final device was fully transparent, enabling real-time optical measurements while selected flow rates were provided by syringe pumps (PhD 2000, Harvard Apparatus, USA).

### Videomicroscopy

*In situ* imaging of cells was conducted using an inverted microscope (IX73, Olympus, Canada) equipped with a 10× Olympus Plan Fluorite objective lens with a 0.30 numerical aperture. Sequential time-lapse imaging was conducted with an 8-second delay for approximately 40 minutes using a CCD camera (Lumenera Infinity 31 U, Ottawa, Canada). Image sequences were then transferred to a computer for analysis, which was carried out using the Fiji distribution of ImageJ software (NIH, US).

### Cell seeding and application of shear stress

*D. discoideum* cells (DH1-10 or *phg2*) were cultured in HL5 medium in 10 cm plastic culture dishes (Falcon) until they covered 60% to 80% of the dish surface. The cells were pelleted by centrifugation, and the resulting cell pellet was resuspended in fresh medium at one-tenth of the initial volume. These suspended cells (the inoculum) were then introduced into a pre-sterilized microfluidic chip, which was pre-filled with HL5 medium. The inoculum was injected via syringe pump set at a near-zero flow rate (0.1 mL h^−1^) until initial cell densities between 100 to 1000 cells mm^−2^ were achieved. Static conditions (zero flow rate) were never applied to avoid nutrient depletion, which can rapidly occur due to the small on-chip volumes. Subsequently, the inoculum was replaced with a culture medium, which was first introduced at a low flow rate (0.1 mL h^−1^) until all culture medium was washed out. Under hydrodynamic conditions of the sterile culture medium, unattached cells were flushed out of the device leaving only the attached cells in the channels. Then, volumetric flow rates of 2, 5, and 10 mL h^−1^ were applied to probe the effect of fluid shear stresses on the cells. The hydraulic retention times were 60, 24, and 12 mins (1, 2.5, and 5 medium recharges per hour) for flow rates of 2, 5, and 10 mL h^−1^, respectively. A new device was used for each cell seeding experiment. Cell counts were manually conducted at each time point. At each time point, the number of detached cells was counted and normalized to the initial count to determine the ratio of remaining cells.

### Statistical analysis

Statistical analyses were conducted using GraphPad Prism version 10.1.2. Data are expressed as the mean ± standard error (σ/n, where σ is the standard deviation and n is the number of samples) of the mean from at least three experiments per condition. Cell densities ranged from 100 to 1000 cells mm^−2^, representing cell densities similar to those during the pre-aggregation phase. We avoided cell densities higher than 1000 cells mm^−2^ to minimize cell-to-cell contact, which can significantly influence adhesion, detachment and motility. Statistical analyses were conducted to ensure significance between data sets that were run with different conditions (e.g., cell types, flow conditions). Differences between any two data sets were evaluated using unpaired t-tests, whereas comparisons among more than two data sets were performed using two-way ANOVA followed by Tukey’s post-hoc test. Statistical significance was defined as a p-value less than 0.05. The levels of significance are denoted in figures as p < 0.05 (*), p < 0.01 (**), p < 0.001 (***), and not significant for p > 0.05 (n.s.).

### Computer simulation

In this study, computational fluid dynamics (CFD) was used to simulate the hydrodynamic flow conditions within a cell-adhesion chamber, with a specific focus on calculating the shear stress exerted on immobilized amoeba cells. The simulation was performed using simulation software (COMSOL Multiphysics 4.2a, Stockholm, Sweden). The simulation employed the fluid-structure interaction to concurrently analyze the effect of fluid flow across the amoeba, under simplified cellular adhesion patterns on the solid surface. The model geometrical model features the microfluidic system, including two inlets and one outlet, and amoebae being represented as solid semi-spherical objects with radius of 5 µm based on approximations from microscope imaging. We fixed the cells at the center of the cell-adhesion chamber (**Figure 1**) to avoid wall effects. A selection of important model properties is presented in **Table 1**.

**Figure 1:**
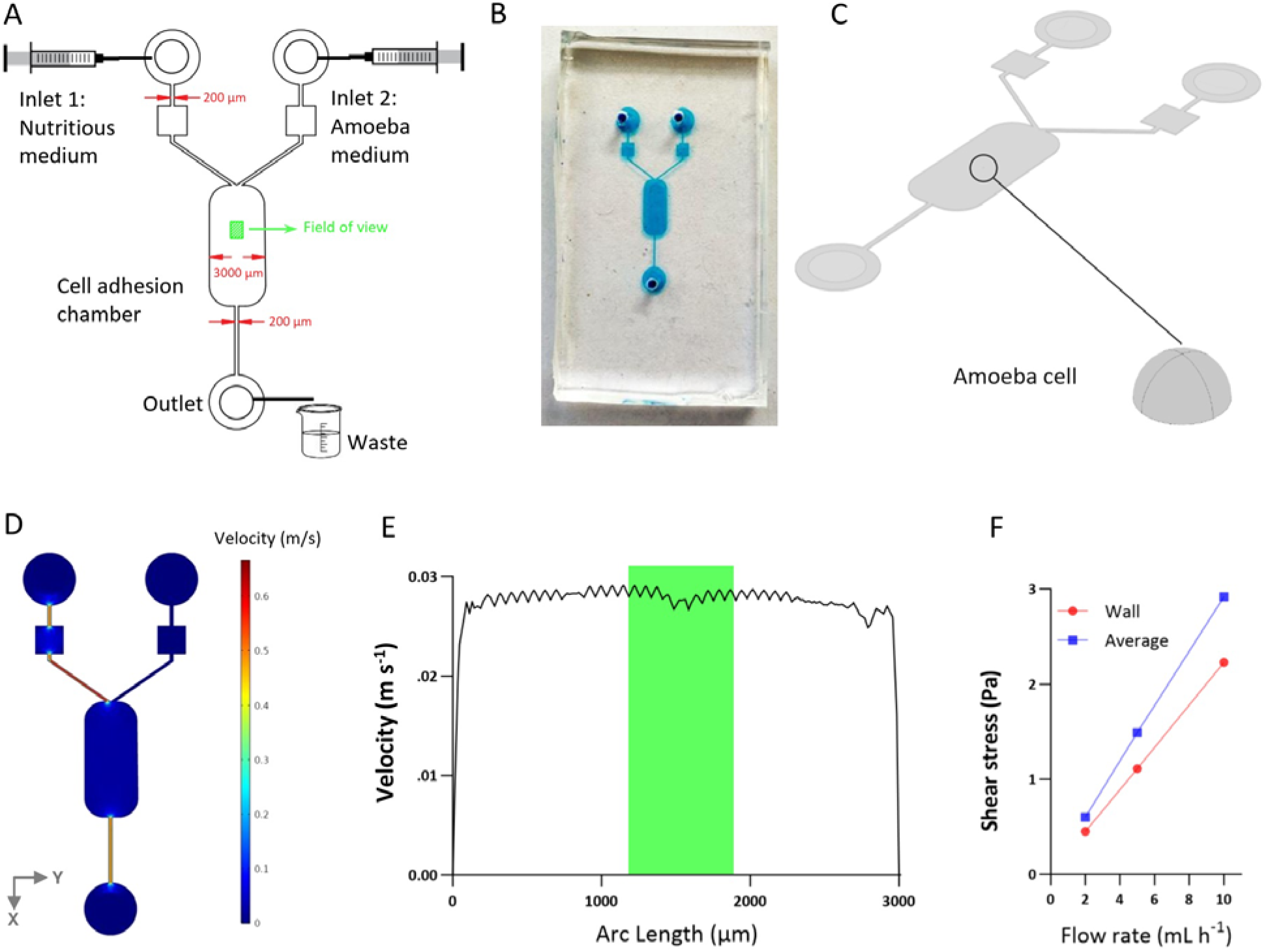
Microfluidic device design. (A) CAD model of the microfluidic array, featuring 2 inlets, 1 outlet, and a cell adhesion chamber, designed to ensure controlled and uniform shear stress, provide ample space for cell mobility, and enable short-term and long-term experiments. (B) Microfluidic chip fabricated based on the CAD design in (a). (C) 3D numerical model of the device with a semi-spherical amoeba model fixed in the middle of the cell adhesion chamber. The shape of the cell model is shown zoomed in for clarity. (D) Velocity magnitude along the device at a flow rate of 5 mL h^−1^ at inlet 1, showing a uniform distribution of velocity across the device. (E) Velocity magnitude along a line through the cell adhesion chamber width (Y-direction), highlighting the uniformity of shear stress distribution; the green box shows the width of the field of view in (A). (F) Difference between the analytically calculated wall shear stress and the numerically calculated average shear stress applied to the amoeba cell surface.

**Table 1:** Properties of the microfluidic device model used in CFD simulation.

|  |  |
| --- | --- |
| **Amoeba radius** | 5 $\mu\text{m}$ |
| **Fluid delivery channel width** | 200 $\mu\text{m}$ |
| **Cell-adhesion chamber length (l)** | 6.6 mm |
| **Cell-adhesion chamber width (w)** | 3 mm |
| **Channel height (h)** | 50 $\mu\text{m}$ |
| **Fluid density** | 1000 $\text{kg m}^{-3}$ |
| **Fluid viscosity ($\mu$)** | 1 mPa s |

The pressure-driven flow of an incompressible liquid through the microfluidic channel can be described using the Navier-Stokes and the continuity equation. To simulate flow through the microfluidic shear stress generator, we used the Poiseuille model to analytically investigate the wall shear stress in a cell-adhesion chamber. The shear stress (τ) is a function of shear rate (γ), which is determined by the volumetric flow rate (Q), the dimensions of the channel height and width (h and w, respectively), and the liquid viscosity (μ):

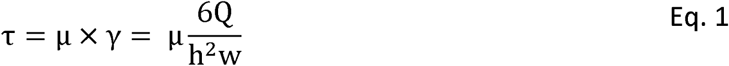

A mesh was developed to define spatial locations where hydrodynamic values and forces were simulated by CFD. A mesh refinement step was conducted iteratively, with enhanced mesh density, until variations in the computed shear stress on the amoeba cell were negligible. A “finer” mesh, resulting in a simulation error of less than 1%, was determined to be optimal **(Table 2)**.

**Table 2:** Summary of mesh independency analysis results.

| **Element size** | **Number of mesh elements** | **Shear stress (Pa)** | **Relative error** |
| --- | --- | --- | --- |
| Coarse | 59442 | 1.87 | 36% |
| Normal | 83001 | 2.59 | 12% |
| Fine | 119616 | 2.91 | 1.2% |
| Finer | 434694 | 2.92 | 0.8% |
| Extra fine | 804595 | 2.95 | - |

## Results

### Device design and shear stress simulation

The design of the final microfluidic shear stress generator, with the design schematic and image shown in **Figure 1A and B**, aimed to meet several key experimental objectives for cell adhesion studies: ensuring a controlled and uniform shear stress across the cell-adhesion chamber, providing ample space for cell motility while minimizing interferences such as wall effect, and enabling measurements at time scales ranging from minutes to more than 12 hours. In this work, all experiments were run for 40 mins. The region of interest (900 × 673 µm) was positioned 1000 µm downstream from the cell-adhesion chamber entry and centered within the cell-adhesion chamber so that the visualized amoeba was not influenced by the vertical sidewalls. We validated this assumption using numerical simulations **(Figure 1C),** demonstrating that along the Y-axis (perpendicular to the flow direction), over 95% of the channel maintained a uniform velocity pattern throughout the channel width, with the exception of locations directly beside the walls **(Figure 1D, E**). This uniformity is due to the low aspect ratio design of the channel (channel height divided by channel width), which minimized the influence of the sidewalls. The low aspect ratio also provided adequate space for cell adhesion, movement, and growth.

According to Eq. 1, the wall shear stresses in the cell-adhesion chamber at flow rates of 2, 5, and 10 mL h^−1^ are calculated as 0.45, 1.11, and 2.23 Pa, respectively. However, due to flow disturbances at the microscale around the cells, the actual shear stress experienced by the cells may differ from the wall shear stress. To assess this effect, we modeled the presence of an idealized cell as a semi-sphere with a radius of 5 µm (**Figure 1C**). The model indicated that the average shear stresses on the cell surface at flow rates of 2, 5, and 10 mL h^−1^ were 0.60, 1.49, and 2.92 Pa, respectively. These values represent 35.5%, 33.4% and 33.13% higher shear stresses experienced by the cell at 2, 5, and 10 mL h^−1^, respectively, compared to the shear stress applied to the channel wall (**Figure 1F**). It is important to note that our simulations did not account for dynamic changes in cell shape. In the future, higher-resolution imaging combined with automated image analysis could be applied to obtain more detailed information about the dimensions on a cell-by-cell basis during the different stages of the experiments and then used to calculate the precise shear stress.

### Cell tracking and migration

The cells exhibited normal migration before detachment. In advance of shear-induced detachment, the cells initially extended several forward and lateral pseudopods. Eventually they ceased migration, became more rounded with less contact to the surface, and then finally detached **(Figure 2A)**. This dynamic response indicates an adaptive behavior to shear stress conditions.

**Figure 2:**
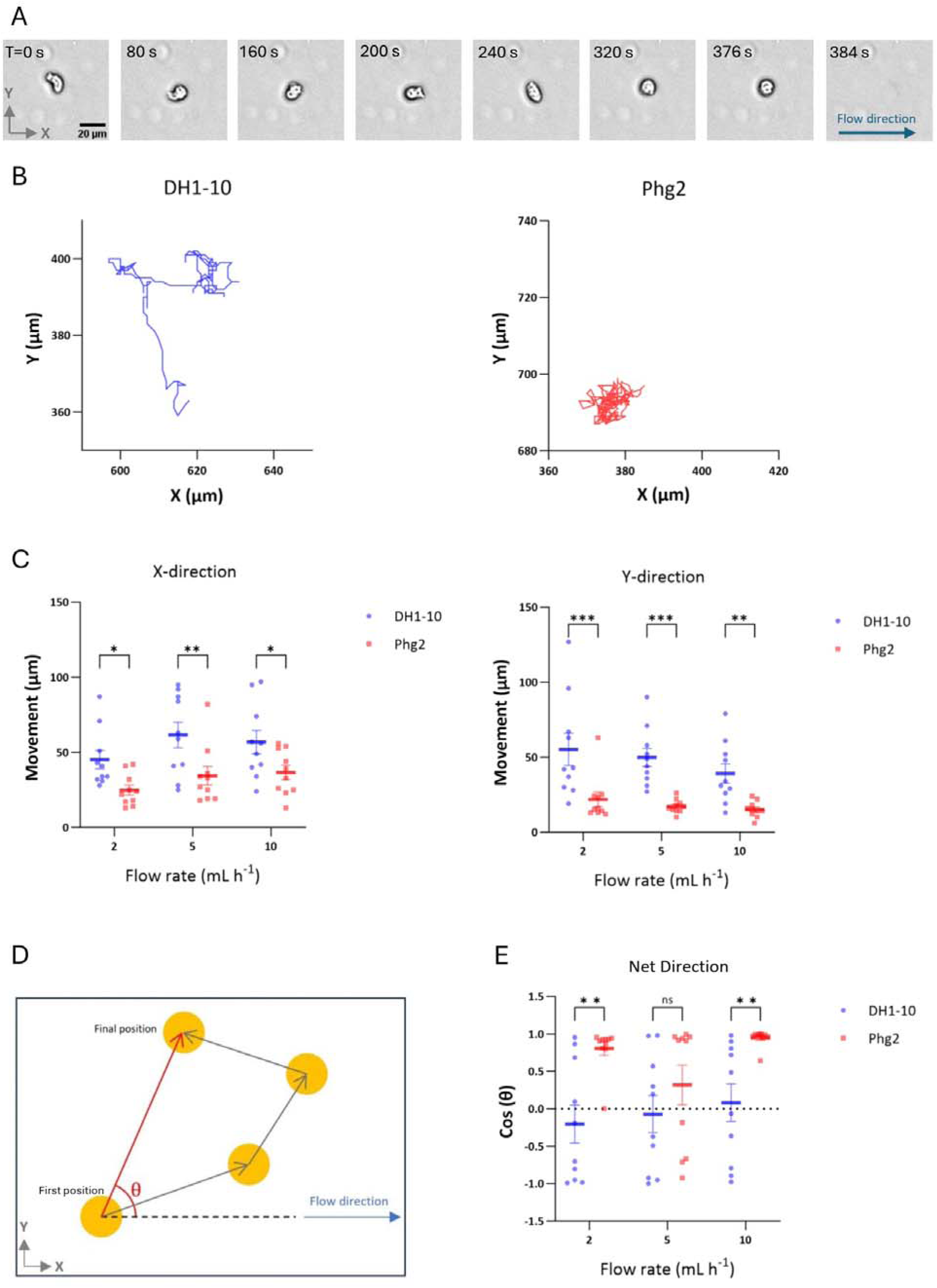
Analysis of *D. discoideum* cell motility under shear stress. (A) Detachment of a single DH1-10 cell at 10X magnification, at 10 mL h^−1^; the arrow in the last frame indicates the flow direction. (B) Migration of a single DH1-10 and single *phg2* cell under fluid flow over a period of 40 minutes. (C) Cell migration of 10 cells under shear stress. (D, E) Mean directionality of cell movement as a function of applied shear stress, indicating to what extent migration is aligned with flow direction. Directionality is defined as the angle between the flow direction and the cell movement direction over a period of 40 min. Therefore, cos(θ)=1 indicates fully biased cell movement in the flow direction, cos(θ)=0 indicates cell movement perpendicular to fluid flow, and cos(θ)=-1 indicates cell movement opposite to fluid flow direction. Error bars represent the standard error.

The motility of the attached cells was quantified by measuring the maximum absolute displacement in the X-direction (parallel to flow) and Y-direction (perpendicular to flow) at each flow rate **(Figure 2B, C)**. DH1-10 cells showed greater movement in both directions compared to *phg2* mutant cells, suggesting higher cellular activity and response to shear stress. When evaluating the effect of flow on motility within the same cell class, the trends indicated that motility in the direction of flow increased and motility perpendicular to flow decreased. However, the statistical differences were lower than our identified threshold for significance.

The directionality of net cell movement was further evaluated by measuring the angle between the initial (t = 0 minutes) and final cell positions (t = 40 minutes), as illustrated in **Figure 2D**. DH1-10 cells exhibited diffusive behavior, characterized by random movement within their environment, and maintained this diffusive pattern even under high shear stress **(Figure 2E)**. In contrast, *phg2* mutant cells demonstrated a movement predominantly aligned with the flow direction (X-direction), as indicated by Cos(θ) values close to 1 **(Figure 2E)** indicating a more pronounced response to the flow conditions. In contrast, movement distance in the X-direction was approximately equivalent for movement upstream compared to movement downstream for DH1-10 cells, indicating a motility that was nearly independent of flow rate.

### Cell detachment under shear flow

We investigated the adhesion and detachment dynamics of *D. discoideum* cells under varying shear stress conditions using the microfluidic shear stress generator. By comparing the detachment and movement of *D. discoideum* wild-type cells and adhesion-defective mutant cells, we aimed to validate the ability of our microfluidic shear stress generator to quantitatively assess comparative cell adhesion.

Various shear stresses were applied by adjusting the inlet flow rate, based on Eq. 1. The flow rate was carefully selected to ensure that both cell movement and detachment could be observed for *D. discoideum* DH1-10 and *phg2* cells. A low flow rate of Q = 2 mL h^−1^ was identified in preliminary assays as the minimum required to study the cell detachment percentage in DH1-10. Below this flow rate, the number of detached cells was found to be insignificant. On the other hand, a high flow rate of Q = 10 mL h^−1^ was sufficient to study cell detachment in DH1-10. This approach allowed us to assess the effects of varying shear stress on cell adhesion and detachment for both cell types. In this study, we were interested in the average results over a range of cell densities from 100 to 1000 cells mm^−2^, a range relevant to the initial densification during the pre-aggregation phase. According to our work, pre-aggregation cell densities can reach 3000 cells mm^−2^ but result in complications related to cell-to-cell contact, which we sought to avoid in this work.

Generally, detachment was initially rapid and slowed until it reached a pseudo-plateau where changes to the number of cells were nearly constant over the time scale of the 40-minute experiment. Cell detachment analysis was conducted, with the results presented as curves in **Figure 3A** and **3B** for DH1-10 and *phg2* amoeba, respectively. The DH1-10 cell type exhibited a gradual increase in detachment percentage and a lower pseudo-plateau in detachment levels across all flow rates compared to that of *phg2*. Based on the two-way ANOVA test, all data were statistically significant, (for example, for DH1-10 detachment at 5 and 10 mL h^−1^). The initial rate of cell detachment and the total percentage of detached cells after reaching a plateau increased with the flow rate but again were notably less affected compared to *phg2* cells, which generally were prone to detachment at all flow rates. In both cases, the detachment process did not reach 100% before the plateau region, indicating that strains contained a population subset with stronger attachment properties. This though this strongly attached population was larger for the DH1-10 cells than for the *phg2* cells. Overall, *phg2* cells have weaker adhesion, including a heightened sensitivity to shear forces.

**Figure 3.**
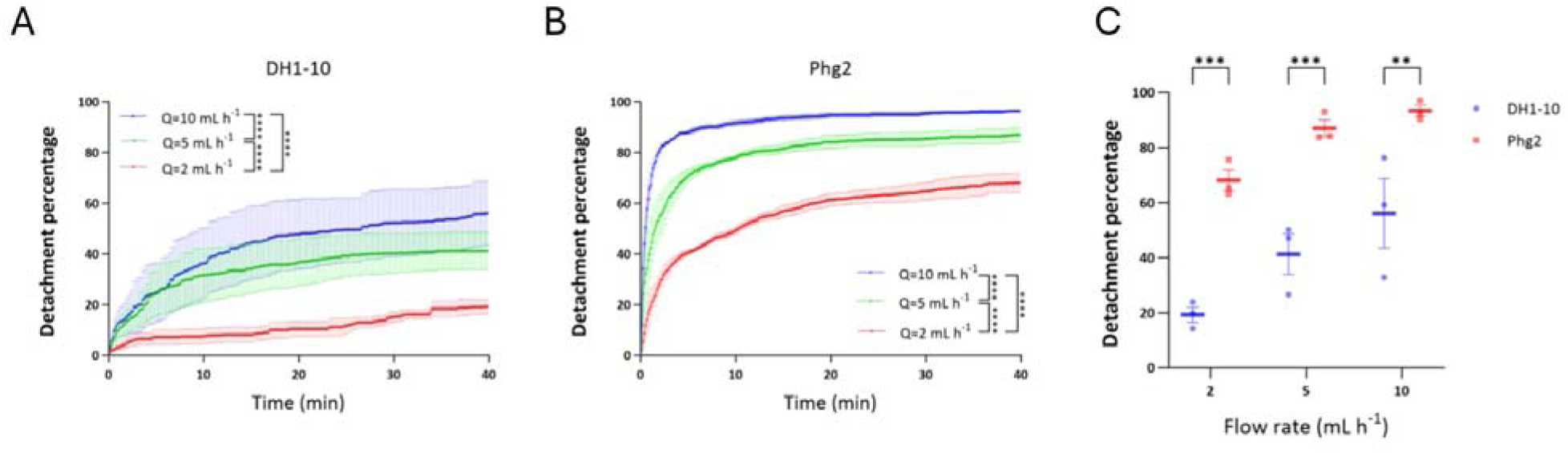
Comparative analysis of cell detachment percentage under various shear stress conditions. (A) Cell detachment curves for *Dictyostelium discoideum* DH1-10 in medium at three flow rates of Q = 10 mL h^−1^ (blue), Q = 5 mL h^−1^ (green), and Q = 2 mL h^−1^ (red), illustrating increased detachment with higher shear stresses from increasing fluid flow rates. (B) Cell detachment curves for the *phg2* adhesion-defective mutant in medium at three flow rates of Q = 10 mL h^−1^ (blue), Q = 5 mL h^−1^ (green), and Q = 2 mL h^−1^ (red), showing a rapid increase in detachment levels. (C) Differential response of DH1-10 and *phg2* final detachment percentages (after 40 mins) for flow rates of 2, 5, and 10 ml h^−1^.

We further analyzed the final detachment percentages of DH1-10 and *phg2* mutants after arriving at the pseudo-plateau (**Figure 3C**); it is evident that *phg2* cells are less adherent and detach more readily in response to fluid flow, which is indicative of weaker adhesion. Conversely, DH1-10 cells display more robust adhesion, demonstrating resistance to shear-induced detachment. These outcomes were anticipated and likely reflect the intrinsic adhesion properties of each cell type.^47^ This differentiation not only highlights the utility of our microfluidic design in assessing cell adhesion under dynamic conditions but also provides valuable insights into the cellular mechanisms governing adhesion, with potential implications for understanding various biological processes and diseases.

### Effect of cell density on detachment

Next, we deepened the analysis by evaluating the effect of initial cell density (number of initial cells per unit area) on the detachment process to test our hypothesis that this may affect the accumulation of cells during the pre-aggregation phase. To begin, we ran simulations to determine how the shear stress was modified on a cell in the presence of a single upstream cell. Using modeling, we accounted for the effect of a single upstream amoeba on the applied shear stress on a (second) neighboring amoeba (downstream). When the intercellular separation distances were large (over 25 µm), the shear stress on the downstream amoeba was the same as reported in Figure 1F (τ = 0.6, 1.49, and 2.92 Pa at 2, 5, and 10 mL h^−1^, respectively). When the separation distances were reduced, our simulation found that the downstream amoeba was partially shielded, resulting in a lower applied shear stress. The results indicated that, at all three flow rates, the presence of an upstream amoeba at an edge-to-edge distance of 10 µm (twice the amoeba’s radius) resulted in a reduction in shear stress on the downstream amoeba of 10%, whereas a 16% reduction in shear stress was observed when the distance was reduced to 5 µm. Next, we extended this simulation to determine the effect of several upstream amoeba at a cell density that mirrored that of our experiments. Based on the initial cell densities in our experiments, we estimated that the average intercellular distance varied significantly, from 7 µm (at 1068 cells mm^−2^) to 55 µm (at 76 cells mm^−2^). We re-ran the simulation with 0, 2, 4, and 7 upstream amoebae, each separated by an edge-to-edge distance d (**Figure 4A**). We conducted these simulations at the three flow rates used in this study and changed the inter-cellular distances to either d=7 or 55 µm to represent the minimum and maximum average cell-to-cell distances based on the range of cell densities in our experiments. As seen in **Figure 4B**, the effect of more upstream amoebae serves to further reduce the applied shear stress on the final amoeba. From these simulations, we see that the shear stress on the most downstream amoeba reaches a stable value after approximately 4 upstream cells for the highest flow rate used. Based on a close analysis, this stability is reached earlier at lower flow rates. Therefore, we are confident that the results with 4 or more upstream amoebae represents an accurate average applied shear stress applied on most of the cells in the experiments. For dense colonies (d=7 µm), the applied shear stress on the most downstream cell was 2.25 Pa at 10 mL h^−1^, marking a reduction of nearly 25% in the applied shear stress compared to that on a single cell. In the case of the lowest cell density (d=55 µm), the absolute shear stress values were nearly unchanged, with only a small reduction from 0.6 Pa to approximately 0.5 Pa after the flow passed by 4 upstream cells with d=7 µm.

**Figure 4.**
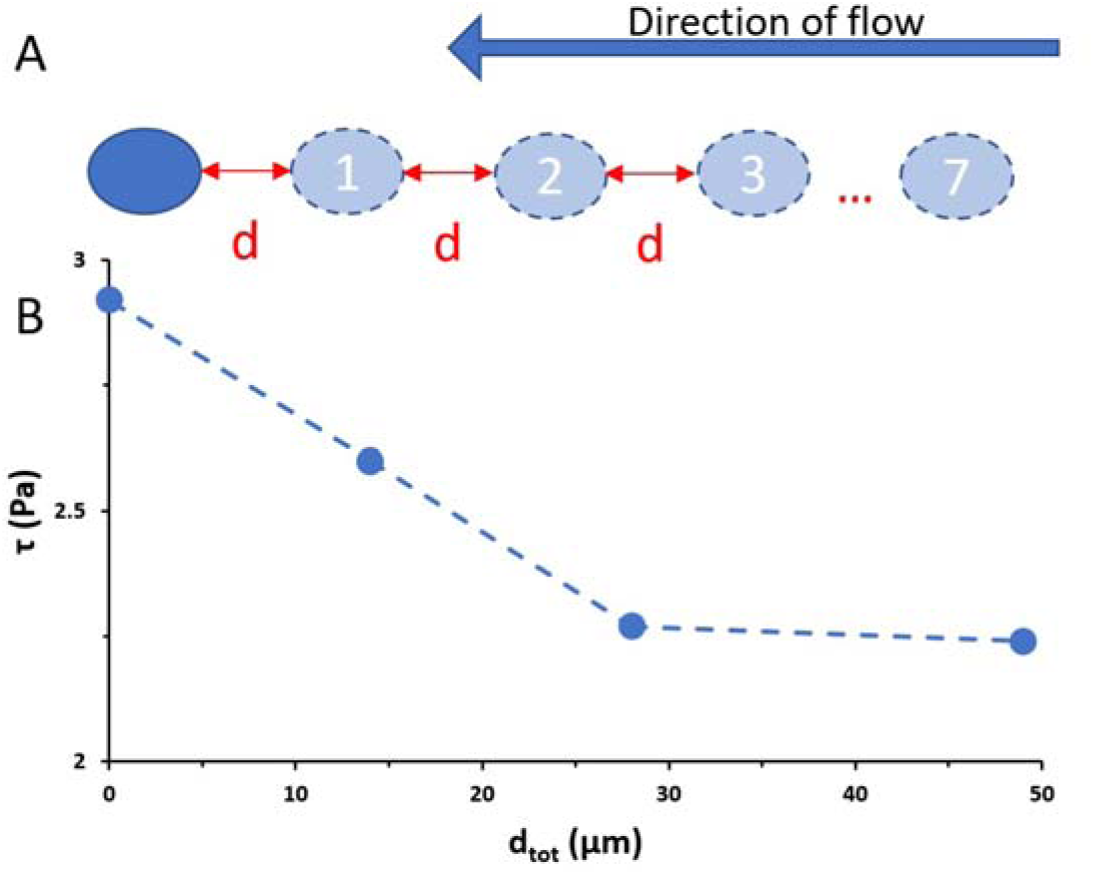
Simulation of shear stresses on an amoeba and accounting for the influence of upstream cells. (A) Schematic of the simulation showing a test cell (dark blue), from which the shear stresses are obtained, and up to 7 upstream cells (light blue) that are separated by a distance d. (B) Shear stress (τ) as a function of the total distance (d_tot_) to the most distant amoeba with data points for number of cells equal to 0, 2, 4, and 7.

Upon investigating the data presented in **Figure 5**, we concluded that cell density can indeed affect detachment, but that this effect is dependent on cell type and flow rate. For example, DH1-10 wild-type cells (**Figures 5A-C**) did show a lower sensitivity to cell detachment when the initial cell densities were high. Unfortunately, the highest cell density for the 2 mL h^−1^ experiment was lower than for the others, so the data have a more pronounced appearance for higher flow rates. In contrast, for *phg2* mutant cells, regardless of the initial cell density, the detachment behavior remained nearly consistent at 2 mL h^−1^, 5 mL h^−1^, and 10 mL h^−1^ (**Figures 5D-F**). This suggests that the reductions in applied shear stresses at high cell densities were not significant enough to impact the very weak adhesion arising from the *phg2* mutation.

**Figure 5:**
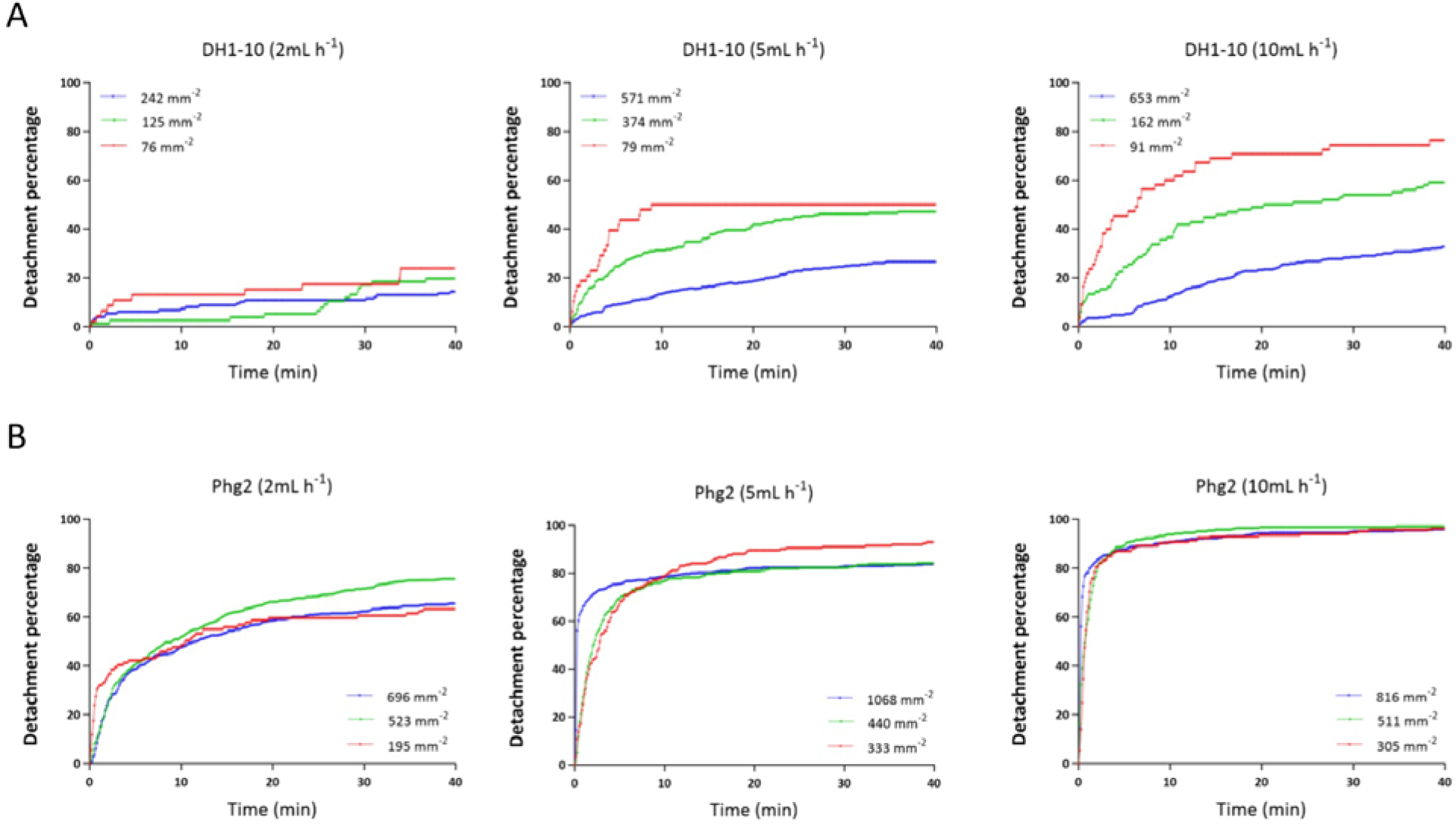
Detachment percentages of (A) DH1-10 wild-type cells and (B) *phg2* mutant cells at different initial cell densities under flow rates of 2 mL h^−1^, 5 mL h^−1^, and 10 mL h^−1^.

## Discussion

Our study employed a custom-designed microfluidic shear stress generator device to analyze the adhesion and detachment of *D. discoideum* DH1-10 and *phg2* mutant cells under various shear stress conditions. The results revealed distinct detachment behaviors between the wild-type DH1-10 and the *phg2* mutant cells. Wild-type DH1-10 cells exhibited a gradual increase in detachment rates with increasing shear stress, whereas *phg2* cells demonstrated a rapid and substantial detachment response. This observation confirms that *phg2* cells possess inherently weaker adhesion properties or an increased sensitivity to shear stress, likely due to the disrupted function of the Phg2 protein. This observation aligns with previous studies, highlighting the crucial role of Phg2 in organizing the actin cytoskeleton, regulating adhesion molecules, and enabling signal transduction for mechanical stimuli response.^47,53^ In *phg2* mutant cells, these processes are disrupted, leading to disorganized actin filaments and altered adhesion properties, which results in increased detachment under shear stress, as our study confirms. Our results on the role of the Phg2 protein in cell adhesion align with those of studies on other adhesion-related proteins such as talin^14,54^ and myosin II.^55,56^ For example, mutants lacking talin have been shown to exhibit weakened adhesion under shear stress, similar to the *phg2* mutant in our study.^54^

In comparison with the literature, our study provides insights into the detachment and cell motility under shear stress conditions. First, our results compare well with those of Décavé et al., who reported cell motility and directionality at higher shear flow rates. Specifically, our observations on the directionality and distance of the cell movement along the flow direction for both DH1-10 and *phg2* cells are coherent with the literature.^14^

A major deviation from the literature, however, are our results showing that cell density plays a measurable effect on the detachment rate of wild type DH1-10 amoeba due to hydrodynamic shielding from neighboring cells, as confirmed by simulations. This is in contrast to the known loss of adhesion that occurs at high cell densities. However, such studies usually have investigated the highest surface coverages (e.g., up to 3000 cell mm^−2^), which result in the destabilizing effects of cell-to-cell contact. In this work, we largely avoided this effect by limiting cell densities to less than 1000 cell mm^−2^.

The use of microfluidic devices in studying *D. discoideum* adhesion under controlled shear stress provides precise control and real-time observation of cell detachment, offering significant advantages over traditional methods. Our approach using large culture areas offers key strengths, including facility in fabrication, accurate application of uniform shear stress over large distances and observation over long durations. Consequently, this design is optimal for investigating dynamic cellular responses and the mechanisms of adhesion and detachment over time. Furthermore, the design enables cells to freely move and thrive within the device, further supporting motility and cell density experiments. Supporting simulations complement the cell density experiments by quantifying the effect of neighbouring cells on applied shear stress, thereby revealing a mutual shielding function that appears to help maintain surface contact in the lead-up to pre-aggregation densification process. Future testing should further develop the model to account for a 2-dimensional cell clusters and more realistic cell shapes, and future experiments should be run under the low nutrient conditions that are usually responsible for triggering aggregation. The versatility of microfluidic fabrication, including various surface functionalizations and architectures, makes our approach adaptable for future studies exploring environmental effects on adhesion, motility and aggregation.

## Conclusion

Our study shows that the microfluidic shear stress generator can be used to effectively assess cell adhesion under varying shear stresses. We used *Dictyostelium discoideum* as a model organism due to its known adhesion properties. Our device measures cell adhesion more accurately than other designs, with minimal wall effects. We compared wild-type DH1-10 cells to adhesion-defective mutant *phg2* cells, finding that the generator distinguishes between cell types based on adhesion. Mutant cells showed weaker adhesion and greater sensitivity to shear forces. The device also allows for detailed analysis of cell adhesion and migration under uniform shear stress, with potential applications in studying mechanotransduction and cell behavior in response to mechanical stimuli.

## Acknowledgements

The authors acknowledge the Natural Sciences and Engineering Research Council (NSERC) for funding under the grant number RGPIN-2020-03603 and the Fonds de Recherche du Québec (FRQ) under the programme Audace. The authors also acknowledge Molly K. Gregas for technical edits.

